# Temperature-dependence and genetic variation in resource acquisition strategies in a model freshwater plant

**DOI:** 10.1101/2023.11.28.569122

**Authors:** Graydon J. Gillies, Amy L. Angert, Takuji Usui

## Abstract

1. Understanding how competition varies with environmental stress is critical to anticipating species and community responses to rapid environmental change. While the stress-gradient hypothesis predicts the strength of competition to decrease with increasing stress, our understanding of how competition varies with stress is limited by a lack of mechanistic understanding of how resource-use traits underlying competitive dynamics respond to stress.
2. Here, we use duckweeds in the *Lemna* species complex to measure how phenotypic and genetic variation in *R\** (a resource acquisition trait representing the minimum resource requirement for positive population growth) varies with high-temperature stress to better understand how stress alters competitive ability for essential resources.
3. We found that heat stress increased the *R\** of *Lemna* plants for nitrogen acquisition. Because lower *R\** values predict dominance in competitive dynamics where resources are limiting, this indicates that under stressful, high temperatures, plants could experience increased sensitivity to competition due to the higher resources required to sustain positive population growth rates.
4. We found minimal genetic variation in *R\** across 11 local genotypes within the *Lemna* species complex, indicating that selection on resource acquisition strategies for essential resources such as nitrogen may be constrained in nature. The expression of genetic variation in *R\** for nitrogen was further reduced under heat stress, suggesting that the response to selection for *R\** could be particularly constrained under high-temperature stress.
5. Contrary to predictions drawn from the gleaner-opportunist trade-off, we did not find evidence for a trade-off in resource acquisition strategies under benign conditions or high-temperature stress. Plants with lower *R\** (i.e., higher growth rates under lower nitrogen levels) were not constrained to have lower growth rates under higher nitrogen levels, possibly because the chosen genotypes have not diverged across resource acquisition strategies or because *Lemna* spp. has escaped this constraint.
6. Importantly, our work outlines that high-temperature stress could increase sensitivity to competition through increased requirement for resources while reducing the evolutionary potential for *Lemna* species to respond to selection for resource traits. This study acts as a key step to understanding the mechanistic traits behind competitive dynamics in resource-limited and stressful environments.

## 1 INTRODUCTION

Competition for limiting resources is a fundamental ecological process that drives patterns of species coexistence (Darwin, 1859; Janzen, 1967; MacArthur et al., 1972). In an era of rapid environmental change, understanding how competitive strengths and outcomes change under novel and stressful environments is of utmost importance. According to the stress-gradient hypothesis, antagonistic interactions such as competition should be stronger in benign environmental conditions and weaker under more stressful conditions (Lortie & Callaway, 2006), possibly due to the greater density, diversity, or per capita effect of competing species under benign environments (Louthan et al., 2015). However, tests of this hypothesis have yielded mixed results in regards to how the strength and outcome of competitive dynamics change across environmental conditions (Hart & Marshall, 2013; Louthan et al., 2015; Malkinson & Tielbörger, 2010). Such inconsistencies in the strength of competition and its interaction with the abiotic environment may potentially be explained by: (1) a more nuanced and mechanistic understanding of how resource-use traits underlying competitive ability are modulated by environmental stress; and (2) understanding how much variation in resource-use traits exists within species and across environments.

In understanding the mechanisms of competition, resource competition theory (RCT) states that individual-level physiological processes can be linked to population-level competitive outcomes by a few, key underlying traits (Tilman, 1982). Notably, a species’ *R\** quantifies the minimum resource level required to sustain positive population growth rates, such that species with a lower *R\** can persist on a lower amount of a given resource (Tilman 1982). Thus, where two species compete for a limiting resource, that with the lower *R\** for that resource may be the superior competitor (Tilman, 1981). Long-standing theory on plant adaptive strategies suggests that under stressful environments, competitive ability may be reduced due to greater diversion of resources towards survival and maintenance (Grace, 1991; Grime, 1977). Returning to the stress-gradient hypothesis, we would predict that weaker competition in stressful environments could be underlain by greater *R\** values in stressful compared to benign environments, due to the greater resource allocation required for population persistence under stressful environments. While previous empirical work has tested for trade-offs between competitive ability and stress tolerance (e.g., Liancourt et al., 2005; Qi et al., 2018), it remains unclear whether differences in competitive ability are mechanistically driven by changes in species’ *R\** values across environments.

Additionally, patterns of competition and resource utilization across stress-gradients could potentially be obscured by large within-species variation in competitive ability or *R\**. Previous studies quantifying the degree of intraspecific variation in competitive ability typically have relied on a phenomenological approach based on demographic models of competition (Adler et al., 2018; Hart et al., 2018) or indirect traits that are proxies for resource allocation to growth (e.g., biomass) and competitive dominance (Johnson et al., 2008; Taylor & Aarssen, 1990). We therefore have much less data on how mechanistic traits that underlie competition for essential resources varies within species, despite such traits providing a direct link from individual- or genotype-level variation in resource requirements to variation in population-level competitive outcomes (Tilman, 1982). Importantly, quantifying genetic variation in *R\** and whether its expression varies across environmental contexts is critical to understand whether in nature, competitive differences can evolve under environmental stress.

While *R\** theory posits that the superior competitor is determined by individuals with lower resource requirements for positive growth, there could also be variation in *R\** if there is a constraint in obtaining low *R\** values, for example, if there is a trade-off between *R\** and maximum growth rate (i.e., gleaner-opportunist trade-offs; Grover, 1990; Stewart & Levin, 1973). Gleaner-opportunist trade-offs describe whether a competitor grows better at low or high resource concentrations. Gleaners are characterized by their ability to grow faster under low resource concentrations relative to their competitors, giving them an edge in nutrient-poor environments. Opportunists have low growth rates in low resource conditions, but exhibit rapid growth when resource concentrations are high. These strategies result in gleaners maintaining a lower *R\** for a given resource while opportunists have higher maximum growth rates on that resource at the cost of a higher *R\** (Grover, 1990). While the importance of such trade-offs in promoting stable coexistence is unclear (Yamamichi & Letten, 2022), quantifying if and how the strength of this trade-off varies with environmental stress could improve our understanding of whether variation in resource acquisition strategies could contribute to the variation in competitive ability we observe across the environmental stress-gradient.

Here, we quantify genetic and environmental variation in *R\** using the widely distributed, freshwater angiosperm, duckweeds (family Lemnaceae) as a model study system (Einhellig et al., 1985). Rapid growth rates and an easily manipulated resource pool makes duckweeds an ideal model species for estimating *R\** values. Furthermore, duckweed populations can easily be cultured under environmental stress, providing insight into whether *R\** may plastically respond to changes in the environment. Specifically, we quantify *R\** across genotypes of common duckweeds in the *Lemna* species complex experiencing benign temperatures or high-temperature stress, with plants competing for nitrogen—an essential resource for photosynthesis and protein synthesis in plants (Novoa & Loomis, 1981). We ask and test a series of questions and predictions:

1. First, how does *R\** vary with benign or stressful temperatures? We predicted that *R\** should be on average higher if plants require greater amounts of resources to persist when grown under high-temperature stress.
2. Second, how much genetic variation in *R\** is there amongst different genotypes in the *Lemna* species complex, and how does the expression of genetic variation vary with heat stress? In other words, is there genetic variation for *R\** such that in nature selection can act on resource traits and does this depend on environmental stress? Previous studies have shown that the expression of genetic variation can differ across environmental contexts, and that stress can limit the expression of genetic variation (Bennington & McGraw, 1996; Hoffmann & Schiffer, 1998) and consequently constrain the response to selection (Emery et al., 2011). For this reason, we predicted that heat-stress should limit the expression of genetic variation for *R\**.
3. Third, are there trade-offs between opportunist and gleaner strategies and does this vary across benign and stressful temperatures? We predicted that in the presence of gleaner-opportunist trade-offs, genotypes that are good competitors at low resource concentrations (i.e., gleaner genotypes with lower *R\**) tend to be worse competitors (i.e., lower maximum growth rate) at high resource concentrations and vice versa. We also predicted that gleaner-opportunist trade-offs could be weaker under heat-stress, if stressful environments increase *R\** while also suppressing maximum growth rates at high resource concentrations.

## 2 METHODS

### 2.1 Species collection and maintenance

Duckweeds are fast becoming a model system for ecology and evolution (Barks & Laird, 2015). Plants are facultatively asexual, with reproduction occurring mostly through rapid clonal budding of daughter fronds (Docauer, 1983). Duckweeds in the *Lemna* species complex can tolerate a wide range of water chemistries and nutrient levels, with a doubling time of 2 to 5 days in optimal conditions (Docauer, 1983; Senevirathna et al., 2023; Ziegler et al., 2015). We sampled 20 unique accessions of the *Lemna* species complex from across 20 sites around Vancouver, British Columbia, Canada, and the Pacific Northwest region including Washington, USA (Table S1). Sampled plants were rinsed under tap water and vortexed with 1X phosphate buffered saline for 5 minutes to remove most surface algae and debris. To obtain axenic cultures of each accession, we then surface-sterilized plants with bleach solution (1% sodium hypochlorite) for 2-4 minutes. We maintained all accessions in the laboratory inside 250 mL Erlenmeyer flasks containing 100 mL of artificial pond media (Appenroth, Teller, and Horn 1996; Table S2). Flasks were placed atop heat mats under ideal temperatures for growth (24 °C) and under LED lighting (SunBlaster 6400K; 16:8 hour light:dark cycle).

We extracted DNA from each accession using a modified CTAB procedure (Healey et al., 2014), barcoded accessions using Tubulin-Based Polymorphism (TBP) fingerprinting for species identification (Braglia et al., 2021), and genotyped each accession using unique microsatellite markers (see Usui & Angert, 2023 for details of extraction and genotyping procedures). Barcoding and genotyping confirmed that our 20 unique accessions were composed of 6 unique *Lemna minor* genotypes and 5 unique genotypes of *Lemna japonica* (Table S1; Usui & Angert, 2023). Both *L. minor* and *L. japonica* currently belong to the morphologically and genetically cryptic *Lemna* species complex. Although some studies have suggested that *L. japonica* is a hybrid between *L. minor* and *Lemna turionifera* (Braglia et al., 2021; Volkova et al., 2023) taxonomic delineation within this species complex is currently unstable. Below, we therefore describe genetic variation in *R\** across 11 unique genotypes in the *Lemna* species complex.

### 2.2 Batch culture experiments

We estimated the minimum nitrogen requirement for positive population growth (i.e., *R\** for nitrogen) in each of the 20 duckweed accessions through batch culture experiments. To obtain an adequate number of plants of each accession and to ensure that each accession was in its growth phase, we first placed plants of each accession into fresh, artificial pond media 8 days prior to the experiment. To minimize the effects of internal nutrient stores obtained prior to the experiment (Docauer, 1983), we then transferred all plants to a nutrient-deficient water medium one day prior to the experiment.

For our batch culture experiment, we then seeded around 10 (SD = 3) individuals of each accession inside a plastic cup (D = 6.6 cm) with 100 mL of artificial pond media and containing varying concentrations of nitrogen (0.01, 0.05, 0.1, 0.5, 1, 5, and 10 mM of nitrate). To impose a temperature treatment, each cup was also placed inside a water bath with temperatures set to 23.4 ± 0.7 ℃ or 36.9 ± 1.3 ℃ (hereafter referred to as ‘benign’ and ‘stressful’ temperatures respectively). These temperatures were chosen as they are close to known estimates of the thermal optimum (*T_opt_*) and critical thermal maximum (*CT_max_*) for *Lemna* spp., respectively (Valentin et al. 2019). Overall, plants of each of the 20 accessions experienced all combinations of nitrogen concentrations (7 levels) and temperature treatment (2 levels) in a fully factorial design. With replication, we therefore tracked population growth rates across 840 experimental populations (N = 3 replicates per accession at each nitrogen and temperature level). These batch culture experiments were performed across 7 temporal blocks (i.e., one nitrogen level per block), with each block having both the benign and stressful temperature treatment, and with the order of nitrogen concentration levels randomized across blocks.

### 2.3 Parameterizing *R\** across temperature and genotypes

To first estimate population growth rates, we counted the total number of individuals in each cup at days 0 and 6 (*N_1_* and *N_2_*, respectively; note that these are individual counts and not nitrogen levels) and estimated relative growth rate (*r*) as:

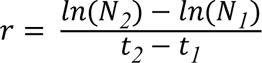

where *t_1_* and *t_2_* are the initial and final days of growth. We then fit a Monod model, which describes the non-linear relationship between growth rates and the limiting resource as:

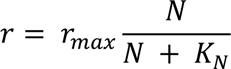

where *r_max_* is the asymptotically-approached maximum growth rate, *N* is the nitrogen concentration, and *K_N_* is the half-saturation constant (i.e., the nitrogen concentration at which growth rate is half of the maximum growth rate). We parameterized Monod models using a Bayesian framework in Stan and R (v4.3.0) using the *brms* package (v2.19; Bürkner, 2017). As *r_max_* and *K_N_* are required to be positive values on the natural scale, we fit these parameters on the log-scale in all models.

To compare resource-dependent growth rates across benign and stressful temperatures and estimate the amount of variance in our parameters explained by genetic variation, we included temperature as a fixed predictor and genotype as a random intercept in the Monod model. We also fit a separate Monod model with an interaction with temperature treatment and genotype as fixed predictors to obtain median posterior estimates of *R\** for each genotype in our experiment across both benign and stressful temperatures (described below). While we use the first model to obtain point estimates at the temperature level and estimate the amount of variance explained by genotype, in the second model we fit genotype as a fixed effect to obtain point estimates at the genotype level, as the use of best linear unbiased predictions (BLUPs) to obtain point estimates from random effects leads to biased estimates and anti-conservative uncertainties (Hadfield et al., 2010). For all models, we used weakly informative normal priors for *r_max_* and *K_N_*, with a mean of –1 and –2, respectively on the log scale (i.e., 0.368 and 0.135 on the natural scale, respectively), and SD of 2. For random effects, we used default half Student’s-*t* priors. We ran models across 4 chains at 5000 iterations each, discarding the first 1000 iterations as warm-up and sampling every 10th iteration thereafter. We assessed for model convergence by visual inspection of traceplots, ensuring no divergence in transitions, and ensuring that chains have reached their equilibrium distribution (as measured by R-hat values ≤ 1).

Using the posterior distributions of *K_N_* and *r_max_* obtained from the Monod models above, we calculated *R\** for nitrogen (i.e., the minimum nitrogen concentration for positive population growth rates) as:

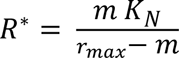

where *m* is the mortality rate (Bernhardt et al., 2020; Tilman, 1976). We estimated *R\** for benign and stressful temperatures, and for each genotype across both temperatures. Since *m* < *r_max_* for an analytical solution to be found in the above equation, we set *m* = 0.01, 0.02 or 0.03 N/day to allow *R\** to be compared across temperature treatment and genotypes. Finally, we tested for the presence of gleaner-opportunist trade-offs in benign and stressful temperatures. We did this by testing for a correlation between *r_max_* and *R\** using the median posterior estimates obtained for each genotype at each temperature level from the Monod model. We used weakly informative normal priors with a mean of 0 and SD of 1, scaled both *r_max_* and *R\** within each temperature level, and then fit a correlation model using *brms*.

## 3 RESULTS

### 3.1 Resource-use efficiency and *R\** across temperature

Monod models showed that the maximum population growth rate (*r_max_*) achieved under stressful temperatures was significantly lower than in benign temperatures (difference in *r_max_* = 0.276, 95% CI: 0.199 to 0.409; Table 2 for model coefficients). In contrast, the half-saturation constant (*K_N_*), which describes the concentration at which growth rates are half of the maximum, was significantly higher in stressful temperatures compared to benign temperatures (difference in *K_N_* = 6.259, 95% CI: 1.649 to 29.874; Table 2). Lower *r_max_* and higher *K_N_* consequently resulted in a shallower growth curve in stressful temperatures (Figure 1A). For a given mortality rate, this means that the *R\** for nitrogen is much greater in stressful than benign temperatures (Figure 1B and 1C), consistent with prediction 1.

**Table 2.** Posterior summaries of maximum growth rate and half-saturation constant (*r_max_* and *K_N_*, respectively). Estimates are from the Monod model of temperature as fixed predictor and with random intercept for genotype. For fixed predictors, we show median estimates and 95% CIs for each parameter on the log scale. Intercept represents estimates at benign temperatures. For random effects, we present mean estimates and 95% CIs of the genotype and residual variance.

| Fixed predictor | Parameter | Median | Lower | Upper |
| --- | --- | --- | --- | --- |
| Intercept (benign) | $r_{max}$ | -1.742 | -1.800 | -1.685 |
| | $K_N$ | -1.926 | -2.211 | -1.646 |
| Temperature treatment (stressful) | $r_{max}$ | -1.287 | -1.612 | -0.893 |
| | $K_N$ | 1.834 | 0.500 | 3.397 |
| Random effect | Parameter | Mean | Lower | Upper |
| Genotype | $r_{max}$ | 0.045 | 0.002 | 0.130 |
| | $K_N$ | 0.283 | 0.019 | 0.701 |
| Residual |  | 0.045 | 0.042 | 0.047 |

**Figure 1.**
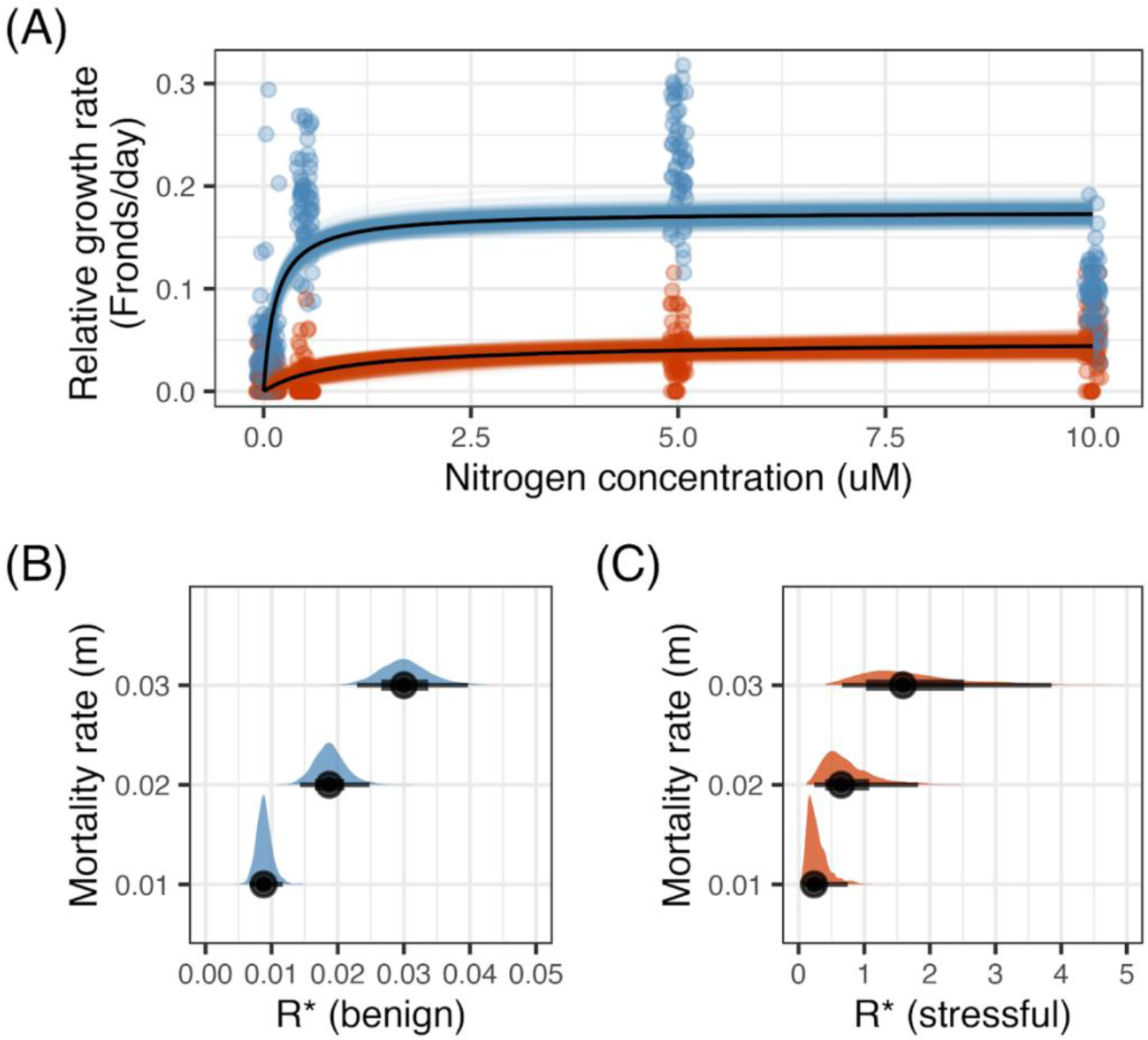
Model predictions on resource-dependent growth rates and *R\** for nitrogen across benign and stressful temperatures. (A) Monod curves describing nitrogen dependent relative growth rates at benign (blue) and stressful (red) temperatures across *Lemna* spp. genotypes. Data points show raw estimates of growth rates (N/day), coloured lines depict uncertainty (i.e., possible posterior draws), and black lines depict median predicted estimates of growth rates at each temperature level. (B and C) Posterior predictions of *R\** across a constant set of mortality rates (*m* = 0.01, 0.02 or 0.03). Histograms represent posterior densities, and circles, thick lines, and thin lines represent the median, 66% and 95% CIs, respectively. Note the difference in scale of x-axes between (B) benign and (C) stressful temperatures.

### 3.2 Resource-use efficiency and *R\** across genotypes

Overall, we detected small amounts of genetic variation in nitrogen-use efficiency across *Lemna* genotypes. Model estimates of genotype variance in *r_max_* was small, with the lower credible intervals close to zero, while genetic variance in *K_N_* was greater by an order of magnitude compared to *r_max_* (Table 2). Median posterior estimates of *r_max_* and *K_N_* across our experimental genotypes confirmed that nitrogen-use efficiency is similar across genotypes (Table S3 and S4). Moreover, posterior estimates of *R\** across experimental genotypes were also similar and with widely overlapping 95% CIs (Figure 2), although estimates were more similar (i.e., more overlap in posterior distributions) across genotypes under high-temperature stress (Figure 2; Figure S2), in line with prediction 2.

**Figure 2.**
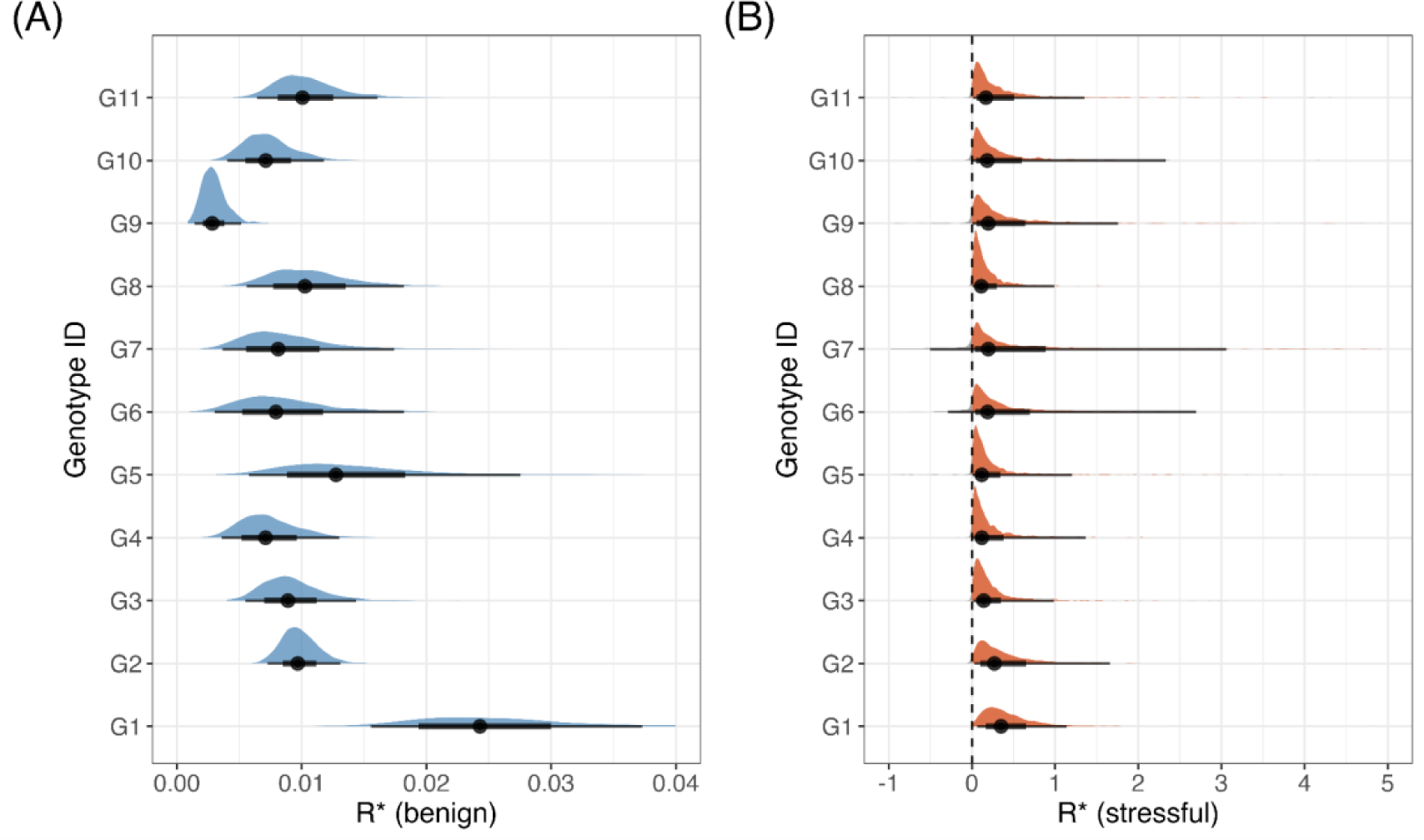
Genetic variation in *Lemna* spp. genotypes for *R\** for nitrogen across (A) benign and (B) stressful temperatures. Histograms represent posterior densities, and circles, thick lines, and thin lines represent the median, 66% and 95% CIs, respectively. The vertical dashed line in (B) indicates *R\** = 0. Note that in stressful temperatures, mortality rates (here set to *m* = 0.01) sometimes exceeded the maximum growth rate achieved, resulting in posterior distributions of *R\** spanning zero (grey shading). These estimates represent conditions where temperatures are too stressful to achieve positive population growth rates regardless of resource availability. Note the difference in scale of x-axes between (A) benign and (B) stressful temperatures. Differences in the overlap of posterior distributions among genotypes are shown in Figure S2.

### 3.3 Gleaner-opportunist trade-offs across temperature

We estimated the degree of correlation between median estimates of *r_max_* and *R\** for nitrogen obtained for each genotype and temperature level (Figure 3). Model predictions show a positive slope between *r_max_* and *R\** in both benign and stressful temperatures under most mortality rates, although the 95% CIs of the slopes are overlapping zero (Table S5). Although the slope between *r_max_* and *R\** became slightly negative under stressful temperatures when *m* = 0.03 (Figure 3C), overall, slopes did not significantly differ among benign and stressful temperatures (Figure 3; Table S5), contrary to prediction 3.

**Figure 3.**
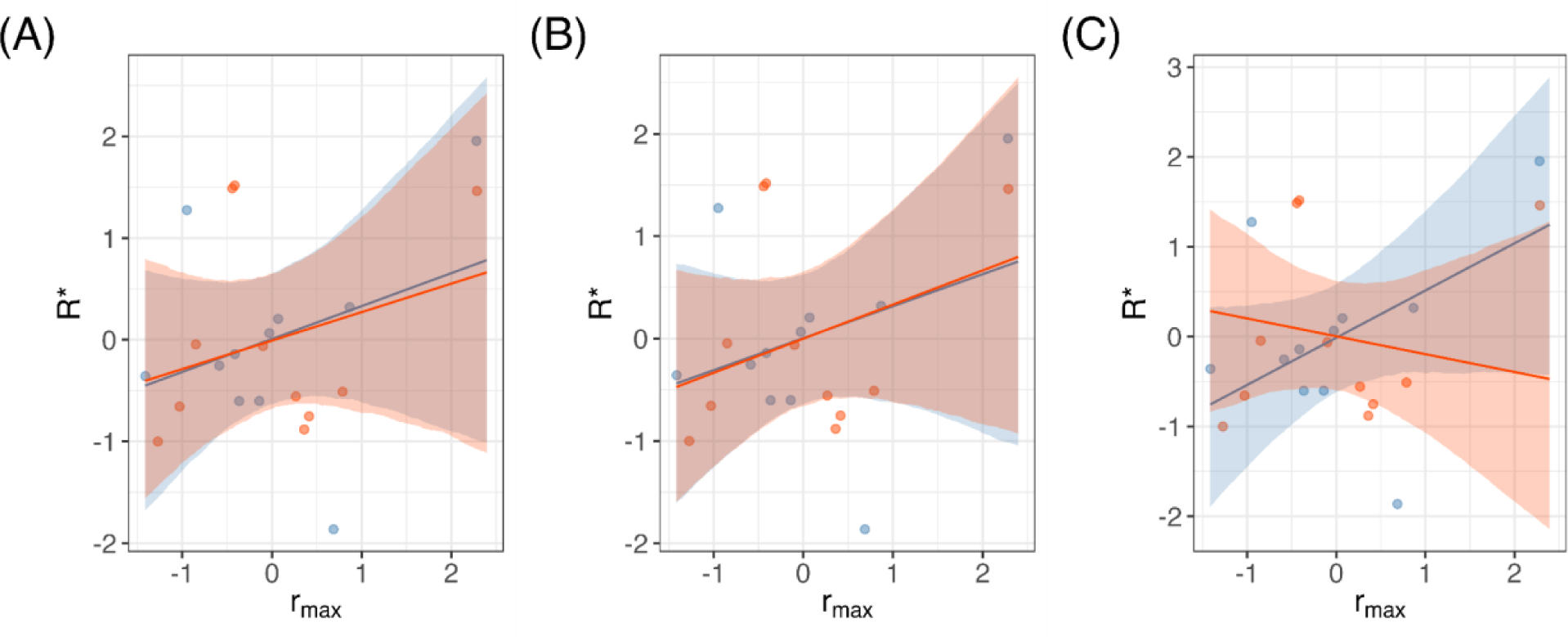
Correlation between *r_max_* and *R\** for nitrogen for benign (blue) and stressful (red) temperatures at mortality rates ranging from: (A) *m* = 0.01; (B) *m* = 0.02; and (C) *m* = 0.03. Data points represent median estimates of *r_max_* and *R\** for each genotype and at each temperature treatment level. Lines and shading represent mean estimates of the correlation coefficient and the 95% CIs. Note that *r_max_* and *R\** have both been scaled within each temperature treatment.

## 4 DISCUSSION

To test mechanistic predictions for how competitive ability differs with environmental stress, we quantified differences in resource-acquisition strategies for an essential resource in terms of the *R\** for nitrogen (i.e., the minimum nitrogen requirement for positive population growth) across a range of nitrogen concentrations and under both benign conditions and high-temperature stress. In line with our prediction, we found that *R\** for nitrogen increases under high-temperature stress, indicating a potential decline in competitive ability due to an increased requirement of this resource to sustain positive population growth under stress. We also found that *R\** varies slightly between genotypes, and consistent with our prediction, we found that the expression of genetic variation for this trait was reduced under stressful temperatures among our experimental genotypes. Lastly, we did not detect the predicted gleaner-opportunist trade-off for resource acquisition in duckweeds under benign or high-temperature stress. In other words, genotypes that had lower R* (i.e., high resource utilization at low resource concentrations) managed to obtain high growth rates at high resource concentrations, and vice versa. Below, we discuss the implications of our results in light of understanding changes to competitive ability under a rapidly changing environment.

### 4.1 High temperature effects on *R*\*

Monod models estimating resource-dependent growth rates and *R\** for nitrogen at both benign and stressful temperatures show that populations exposed to high, stressful temperatures had higher *R\** values than those grown in benign temperatures (Figure 1). This is consistent with the hypothesis that physiological stress caused by exposure to high temperatures necessitates an increase in essential resource requirements to maintain positive population growth rates. While it may be argued that in nature *R\** may be greater due to potentially higher mortality rates in stressful compared to benign environments, our results show that *R\** is greater under heat stress even if mortality rates are held constant between our temperature treatments. This suggests a large effect of temperature stress on *R\**, and that the greater resource requirements to maintain positive growth rates may contribute to a decreased competitive ability under stressful conditions (Tilman, 1981).

In an era of rapid environmental change, understanding how competitive dynamics change with heat stress is important for predicting how the interactive effects of environmental change and species interactions will shape future communities. This will be particularly important as responses to climate change vary within and across species, leading to asynchronous poleward and elevational range shifts and consequent changes in species interactions within novel and existing communities (Alexander et al., 2015). Our work supports the prediction that heat stress causes species to have greater *R\** and therefore increased resource requirements for maintaining population growth, implying that organisms with increased *R\** will need to deal with the dual stresses of both increasing temperature warming and greater sensitivity to competition. Thus, it will be important for future studies to test how resource-use traits respond to climate change in natural populations and understand how these mechanistic traits drive population dynamics in response to both abiotic and biotic change.

Furthermore, there is some evidence suggesting that how the strength of competition varies with environmental stress depends on the different forms of competition. For example, a meta-analysis found that intraspecific root-root competition is generally higher under drought conditions, but that shoot-shoot competition is higher with increased water availability (Foxx & Fort, 2019). Across species, there is also evidence of greater shoot-shoot interspecific competition in marsh plants when nutrient concentrations are high, while root-root competition is predominant in low nutrient concentrations (Emery et al., 2001). With duckweed, while we expect competition for nitrogen to decrease with high nitrogen levels, increased resources could also lead to greater crowding and frond overlap, leading to increased competition for space and light (Docauer, 1983). Interestingly, growth rates under benign temperatures appeared to be lower at the highest nitrogen level compared to more moderate concentrations (Figure 1A). While it is unclear why this pattern was observed, it is possible that overcrowding or competition for light limited growth at the highest nitrogen levels, or that overfertilization resulted in decreased reproductive output, similar to observed decreases in crop yield and quality after overfertilization in agricultural systems (Albornoz, 2016; Cerrato & Blackmer, 1990; Fernández-Escobar et al., 2006). While this could affect the fit of the Monod curve to our data, this did not affect our main conclusions as further sensitivity analysis which removed the highest nitrogen concentration did not qualitatively change our results that heat stress increased *R\** (Figure S1; Table S6). Overall, for greater understanding of how competition changes across environments, future studies will need to consider how different forms of competition, as well as competition for multiple essential resources, vary across environmental stress-gradients.

### 4.2 Genetic variation in *R*\*

Monod models at the genotype level showed that there was a small amount of genetic variation in *R\** across closely related genotypes sampled across local scales (Figure 2; Table 2). Small amounts of genetic variation in the *Lemna* species complex are potentially adequate for responding to selection on resource acquisition strategies in nature. *Lemna* spp.’s exceptionally short doubling time of just a few days (Ziegler et al., 2015) means that these populations could evolve rapidly in response to environmental change through the sorting of genotypes with different resource-use traits. The variation in *R\** between genotypes that we find is consistent with previous work that has found that experimental duckweed populations can evolve rapidly in response to interspecific competition to facilitate coexistence (Hart et al., 2019). Similarly, Bernhardt et al. (2020) demonstrated rapid evolution of *R\** in experimental populations of green algae (*Chlamydomonas reinhardtii*) in response to different resource limitations.

Notably, this genetic variation in *R\** across genotypes was greater under benign than stressful temperatures (see non-overlapping posterior distributions for some genotypes under benign but not stressful temperatures in Figure 2, and differences in posterior distributions among genotypes in Figure S2). This is consistent with previous studies that have shown that environmental stress can limit the expression of genetic variation (Bennington & McGraw, 1996; Hoffmann & Schiffer, 1998) and consequently constrain the response to selection (Emery et al., 2011). In terms of resource use and high-temperature stress, while populations become more sensitive to competition through plastic responses that increase resource requirements for positive population growth, these populations may also be constrained in their genetic responses to selection imposed by competition under heat stress. If true, then this may potentially limit the capacity of populations to adaptively evolve in response to competition under climate change. To assess these processes outside of laboratory systems, future studies should assess how *R\** evolves in natural field populations and how the selection for this trait depends on environmental variation across time and space.

### 4.3 Gleaner-opportunist trade-offs

We found a weakly positive but statistically non-significant correlation between *R\** and asymptotic maximum growth rate (*r_max_*) under both benign and stressful temperatures (Figure 3). The relationship became weakly negative at high temperatures and high mortality rates (*m* = 0.03), although this difference was still statistically non-significant. This result contrasts with the predictions drawn from the gleaner-opportunist trade-off (Grover, 1990, 1997). In other words, for duckweeds in our experiment, we did not find evidence for constrained resource use strategies, and therefore plants that grow faster under lower resource concentrations could also grow fast under higher resource environments.

We may have not observed the predicted gleaner-opportunist trade-off in the *Lemna* species complex because the experimental genotypes were collected at a relatively small scale in the Pacific Northwest (Table S1 in Supporting Information). This somewhat narrow spatial scale relative to the dispersal capacity of duckweed (Coughlan et al., 2015; Les et al., 2003; Silva et al., 2018) may have restricted the variety of environmental conditions experienced by the collected genotypes and consequently limited our ability to collect genotypes that have diverged in resource strategies across the gleaner-opportunist trade-off. Furthermore, if the experimental genotypes were too genetically similar (i.e., have not yet diverged substantially) for different resource strategies to have evolved, we would not detect any trade-off in resource acquisition strategies. This is consistent with our data showing little genetic variation in *R\** across our experimental genotypes. High genetic similarity may be present in the *Lemna* species complex in this region due to the facultatively clonal nature of duckweeds and extremely low mutation rates (Sandler et al., 2020). However, Senevirathna et al. (2023) demonstrated genetic divergence across *L. minor* plants from nearby populations with potential differentiation in water chemistry profiles. It remains unclear whether our *Lemna* spp. genotypes collected from the Pacific Northwest have undergone adequate divergence to evolve resource strategies across a gleaner-opportunist trade-off.

It is possible, however, that we did not observe this trade-off because the *Lemna* species complex as a whole is not constrained by the gleaner-opportunist trade-off. For example, some work has shown that cryptophytes (a phylum of phytoplankton) are not constrained by this trade-off for photosynthetic light acquisition (Swanson et al., 2023). Additionally, a meta-analysis showed that the gleaner-opportunist trade-off does not apply across taxa of heterotrophic eukaryotes (Kiørboe & Thomas, 2020). Perhaps the *Lemna* species complex has, similarly, ‘escaped’ the constraints of this trade-off or are better modelled by an entirely different resource use framework.

## 5 CONCLUSION

Our study demonstrates that plants in the *Lemna* species complex have: (1) higher *R\** values under stressful conditions; (2) limited but measurable genetic variation in *R\** across genotypes; and (3) no detectable gleaner-opportunist trade-offs for nitrogen acquisition. To our knowledge, this work represents the first attempt to quantify changes in *R\** across distinct genotypes when organisms are exposed to stressful environmental conditions. Applied in a broader context, our results have important implications for species responses to a rapidly changing environment. As anthropogenic stressors such as climate warming or habitat degradation accelerate, it is possible that competitive sensitivity will increase due to greater resource requirements to sustain positive population growth rates, leading to cascading ecological effects at the community level. More work is needed to understand how competition and other species interactions will change in response to environmental changes, but this work acts as a first step to quantify species mechanistic responses to heat stress.

