## Supporting Information for "Temperature-dependence and genetic variation in resource acquisition strategies in a model freshwater plant"

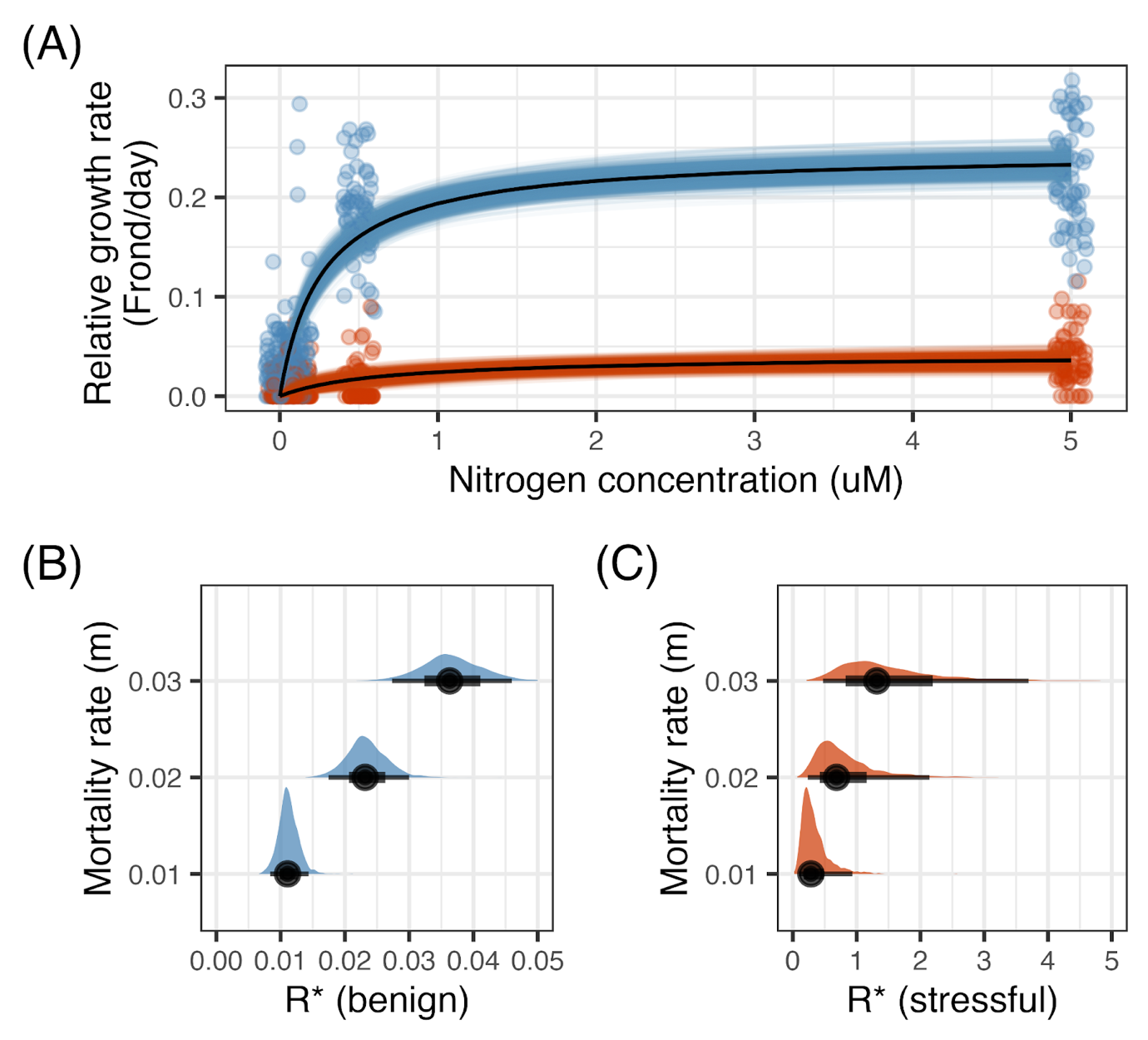

**Figure S1.** Sensitivity analysis of resource-dependent growth rates and *R** for nitrogen across benign and stressful temperatures. Sensitivity analysis was conducted without the highest concentration of nitrogen (10uM). (A) Monod curves describing nitrogen dependent growth rates at benign (blue) and stressful (red) temperatures. Data points show raw estimates of growth rates (N/day), coloured lines depict uncertainty (i.e., possible posterior draws), and black lines depict median predicted estimates of growth rates. (B and C) Posterior predictions of *R** across a constant set of mortality rates (*m* = 0.01, 0.02 or 0.03). Histograms represent posterior densities, and circles, thick lines, and thin lines represent the median, 66% and 95% CIs, respectively. Note the difference in scale of x-axes between (B) and (C).

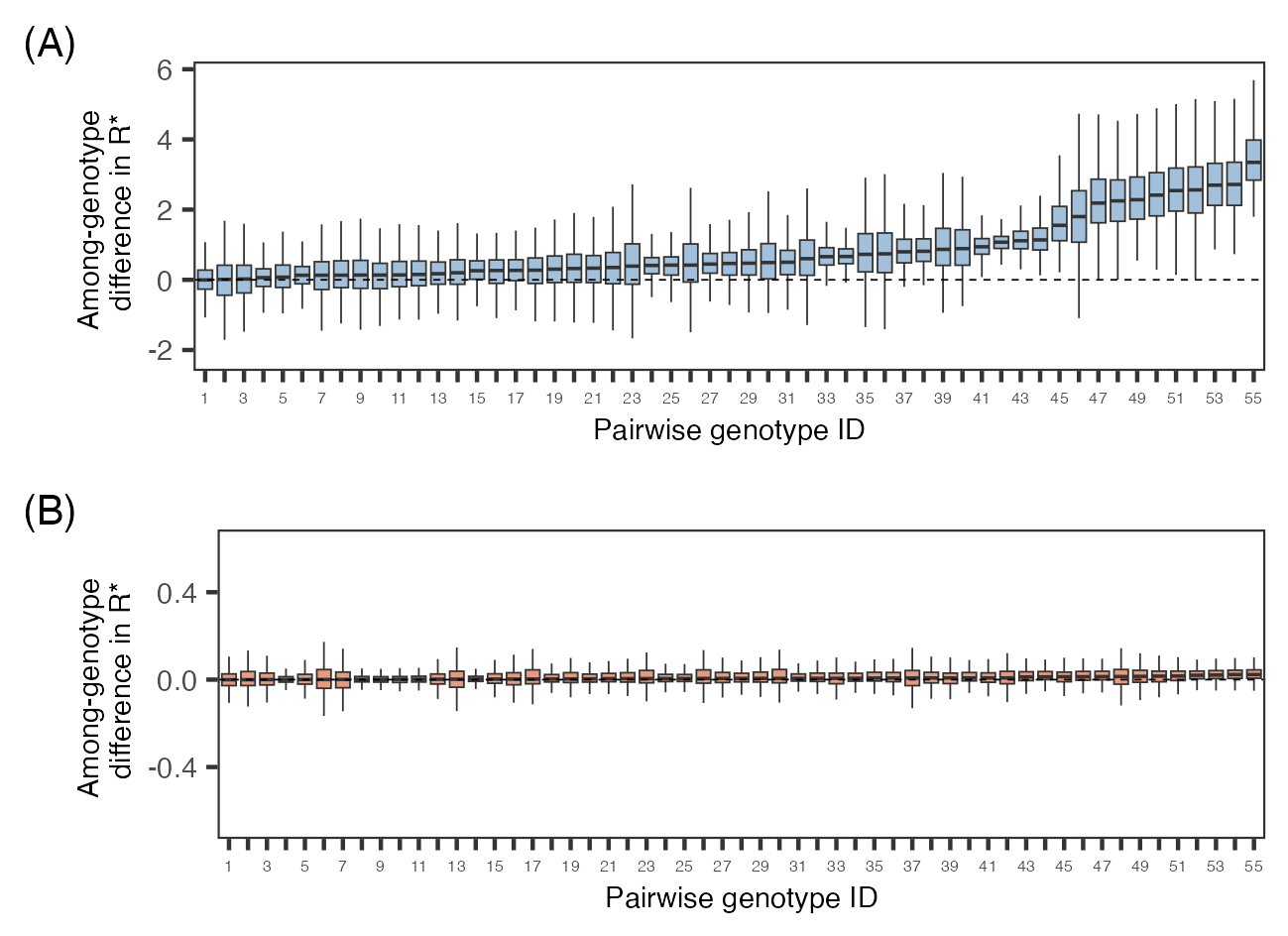

**Figure S2.** Difference in *R** for nitrogen among all pairs of 11 *Lemna* genotypes under (A; blue) benign and (B; red) high-temperature stress. Difference in *R** among genotypes was estimated as the among-genotype difference in the posterior distributions of *R** between each pair of genotypes (all 55 pairwise combinations of 11 genotypes compared under each temperature level), where 0 = no difference and more positive values translate to greater differences between genotypes. Boxplots (representing the median, 25th, and 75th quartiles) are ordered from smallest to largest difference in *R** between a pair of genotypes.

**Table S1.** Sampling locations of experimental *Lemna* plants, where each row represents a unique accession and its site information. All samples were collected between March and November in 2018 and 2019. Species identification was based on TBP barcodes and genotype identification was based on microsatellite markers (see Usui & Angert 2023).

| Species | Genotype ID | Lat | Long | Location |
| --- | --- | --- | --- | --- |
| *L. minor* | G1 | 49.2639 | –123.2499 | Biodiversity Museum, Vancouver, BC |
| *L. minor* | G2 | 49.2564 | –122.9648 | Burnaby Lake (Ditch), Burnaby, BC |
| *L. minor* | G2 | 49.2565 | –122.9652 | Burnaby Lake (West), Burnaby, BC |
| *L. minor* | G2 | 49.2175 | –123.1766 | Celtic Ave & Balaclava St, Vancouver, BC |
| *L. minor* | G2 | 49.2184 | –123.1788 | Celtic Ave & Blenheim St, Vancouver, BC |
| *L. minor* | G2 | 48.6975 | –122.4776 | Hemlock Trail, Bellingham, WA |
| *L. minor* | G2 | 49.1738 | –123.1976 | Terra Nova Park (North), Richmond, BC |
| *L. minor* | G2 | 49.2230 | –123.1762 | W53 Ave & Balaclava St, Vancouver, BC |
| *L. japonica* | G3 | 49.2715 | –123.1107 | Hinge Park (North), Vancouver, BC |
| *L. japonica* | G3 | 49.2706 | –123.1103 | Hinge Park (South), Vancouver, BC |
| *L. japonica* | G4 | 49.2695 | –123.1358 | Granville Island, Vancouver, BC |
| *L. minor* | G5 | 49.6952 | –124.5076 | Cranby Lake, Texada Island, BC |
| *L. minor* | G6 | 48.7366 | –122.3820 | Geneva Pond, Bellingham, WA |
| *L. minor* | G7 | 49.2658 | –123.2599 | Nitobe Memorial Garden, Vancouver, BC |
| *L. japonica* | G8 | 49.2243 | –123.1759 | Southlands Heritage Farm, Vancouver, BC |
| *L. japonica* | G9 | 49.2565 | –122.9647 | Burnaby Lake (North), Burnaby, BC |
| *L. minor* | G10 | 49.2522 | –122.9625 | Burnaby Lake (East), Burnaby, BC |
| *L. minor* | G10 | 49.1722 | –123.1984 | Terra Nova Park (South), Richmond, BC |
| *L. japonica* | G11 | 49.2382 | –122.9705 | Deer Lake (East), Vancouver, BC |
| *L. japonica* | G11 | 49.2383 | –122.9713 | Deer Lake (North), Vancouver, BC |

**Table S2.** Recipe for artificial pond media based on Appenroth et al. (1996). The experimental pond media is made of 4 stock solutions with chemical ingredients listed below. Final concentration is the concentration of each chemical ingredient in the final experimental media after combining all of the stocks with DI water.

| Stock | Chemical ingredients  (mm; g/mol) | Stock conc. | Amount (g/L) | Final conc. |
| --- | --- | --- | --- | --- |
| 1 | KH_2_PO_4_ (136.1) | 30 mM | 4.083 | 0.15 mM |
| 2 | Ca(NO_3_)_2_ 4H_2_O (236.2) | 0.2 M | 47.23 | 1 mM |
| 3 | KNO_3_ (101.1) | 1.6 M | 161.8 | 8 mM |
| 3 | H_3_BO_3_ (61.83) | 1 mM | 0.0618 | 5 $\mu$M |
| 3 | MnCl_2_ 4H_2_O (197.9) | 2.6 mM | 0.5145 | 13 $\mu$M |
| 3 | Na_2_MoO_4_ 2H_2_O (241.95) | 80 $\mu$M | 0.0194 | 0.4 $\mu$M |
| 3 | MgSO_4_ 7H_2_O (246.48) | 0.2 M | 49.30 | 1 mM |
| 4 | FeNaEDTA (367.1) | 5 mM | 1.835 | 25 $\mu$M |

**Table S3.** Posterior summaries of maximum growth rate (*r_max_*) for each genotype ID at each temperature level. Estimates are from the Monod model of temperature and its interaction with genotype ID as fixed predictors. Median estimates and 95% CIs are shown on the log-scale.

| Temperature | Genotype ID | Median | Lower | Upper |
| --- | --- | --- | --- | --- |
| Benign | G1 | –1.485 | –1.639 | –1.335 |
|  | G2 | –1.747 | –2.079 | –1.418 |
|  | G3 | –1.797 | –2.164 | –1.439 |
|  | G4 | –1.763 | –2.173 | –1.372 |
|  | G5 | –1.877 | –2.325 | –1.446 |
|  | G6 | –1.947 | –2.392 | –1.539 |
|  | G7 | –1.825 | –2.267 | –1.431 |
|  | G8 | –1.640 | –2.036 | –1.256 |
|  | G9 | –1.659 | –2.036 | –1.304 |
|  | G10 | –1.792 | –2.158 | –1.429 |
|  | G11 | –1.734 | –2.105 | –1.380 |
| Stressful | G1 | –2.489 | –3.194 | –1.761 |
|  | G2 | –3.298 | –4.659 | –1.903 |
|  | G3 | –2.906 | –4.546 | –1.411 |
|  | G4 | –3.088 | –4.972 | –1.497 |
|  | G5 | –3.043 | –5.207 | –1.378 |
|  | G6 | –3.685 | –5.957 | –1.837 |
|  | G7 | –3.844 | –6.117 | –1.947 |
|  | G8 | –3.049 | –4.781 | –1.476 |
|  | G9 | –3.337 | –5.358 | –1.751 |
|  | G10 | –3.542 | –5.432 | –1.906 |
|  | G11 | –3.196 | –4.952 | –1.653 |

**Table S4.** Posterior summaries of half-saturation constant (*K_N_*) for each genotype ID at each temperature level. Estimates are from the Monod model of temperature and its interaction with genotype ID as fixed predictors. Median estimates and 95% CIs are shown on the log-scale.

| Temperature | Genotype ID | Median | Lower | Upper |
| --- | --- | --- | --- | --- |
| Benign | G1 | –0.634 | –1.132 | –0.136 |
|  | G2 | –1.834 | –2.917 | –0.741 |
|  | G3 | –1.979 | –3.206 | –0.772 |
|  | G4 | –2.176 | –3.510 | –0.891 |
|  | G5 | –1.699 | –3.181 | –0.244 |
|  | G6 | –2.266 | –3.833 | –0.782 |
|  | G7 | –2.111 | –3.589 | –0.679 |
|  | G8 | –1.675 | –2.975 | –0.413 |
|  | G9 | –2.987 | –4.341 | –1.677 |
|  | G10 | –2.203 | –3.469 | –0.989 |
|  | G11 | –1.785 | –2.992 | –0.591 |
| Stressful | G1 | 0.866 | –1.590 | 2.844 |
|  | G2 | –0.424 | –4.789 | 3.401 |
|  | G3 | –0.530 | –4.781 | 3.229 |
|  | G4 | –0.905 | –5.254 | 2.961 |
|  | G5 | –0.851 | –5.228 | 2.931 |
|  | G6 | –1.093 | –5.452 | 2.748 |
|  | G7 | –1.117 | –5.577 | 2.761 |
|  | G8 | –0.910 | –5.203 | 2.902 |
|  | G9 | –0.734 | –5.321 | 3.139 |
|  | G10 | –1.059 | –5.431 | 2.922 |
|  | G11 | –0.701 | –5.060 | 3.148 |

**Table S5.** Model coefficients testing the correlation between *N** and *r_max_* for a range of mortality rates (*m* = 0.01 to 0.03) and for benign and stressful temperatures. The slopes and 95% LCIs and UCIs are given.

| Mortality rate (*m*) | Temperature level | Slope | 95% LCI | 95% UCI |
| --- | --- | --- | --- | --- |
| 0.001 | Benign | 0.327 | -0.362 | 1.006 |
|  | Stressful | 0.287 | -1.337 | 1.921 |
| 0.002 | Benign | 0.301 | -0.367 | 0.974 |
|  | Stressful | 0.350 | -1.261 | 1.966 |
| 0.003 | Benign | 0.249 | -0.428 | 0.900 |
|  | Stressful | -0.198 | -1.810 | 1.452 |

**Table S6.** Sensitivity analysis without the highest concentration of nitrogen (10uM). Estimates are from the Monod model of temperature as fixed predictor and with random intercept for genotype. For fixed predictors, we show median posterior and 95% CIs for each parameter on the log scale. Intercept represents estimates at benign temperatures.

| Fixed predictor | Parameter | Median | Lower | Upper |
| --- | --- | --- | --- | --- |
| Intercept (benign) | *r_max_* | –1.406 | –1.476 | –1.341 |
|  | *K_N_* | –1.343 | –1.632 | –1.070 |
| Temperature treatment (stressful) | *r_max_* | –1.781 | –2.178 | –1.387 |
|  | *K_N_* | 0.931 | –0.389 | 2.223 |
